# A Human Accelerated Region is a Leydig cell *GLI2* Enhancer that Affects Male-Typical Behavior

**DOI:** 10.1101/2021.01.27.428524

**Authors:** Andrew R. Norman, Ann H. Ryu, Kirsty Jamieson, Sean Thomas, Yin Shen, Nadav Ahituv, Katherine S. Pollard, Jeremy F. Reiter

**Affiliations:** Department of Biochemistry and Biophysics, Cardiovascular Research Institute, University of California San Francisco, San Francisco, CA, USA; Department of Bioengineering and Therapeutic Sciences, University of California, San Francisco, San Francisco, CA, USA; Department of Neurology, University of California, San Francisco, San Francisco, CA, USA; Altius Institute for Biomedical Sciences, Washington, USA; Institute for Human Genetics, University of California, San Francisco, San Francisco, CA, USA; Gladstone Institute of Data Science & Biotechnology, San Francisco, CA; Department of Epidemiology & Biostatistics, University of California, San Francisco, CA; Chan Zuckerberg Biohub, San Francisco, CA 94158, USA

## Abstract

Human accelerated regions (HARs) are sequences that have evolved at an accelerated rate in the human lineage. Some HARs are developmental enhancers. We used a massively parallel reporter assay (MPRA) to identify HARs with enhancer activity in a mammalian testis cell line. A subset of HARs exhibited differential activity between the human and chimpanzee orthologs, representing candidates for underlying unique human male reproductive biology. We further characterized one of these candidate testis enhancers, 2xHAR.238. CRISPR/Cas9-mediated deletion in a testis cell line and mice revealed that 2xHAR.238 enhances expression of *Gli2*, encoding a Hedgehog pathway effector, in testis Leydig cells. 4C-seq revealed that 2xHAR.238 contacts the *Gli2* promoter, consistent with enhancer function. In adult male mice, deletion of 2xHAR.238 disrupted mouse male-typical behavior and male interest in female odor. Combined, our work identifies a HAR that promotes the expression of *Gli2* in Leydig cells and may have contributed to the evolution of human male reproductive biology.

## INTRODUCTION

Enhancers regulate developmental processes and mutations in enhancers can result in phenotypic changes subject to evolutionary selection (*1–4*). Demonstrating a causative link between enhancer molecular evolution and phenotypic changes is a challenge, given that most phenotypes are the result of multiple genes, most genes are regulated by multiple enhancers, and the effect of one enhancer on gene activity can be subtle. Conservation of enhancer sequence across multiple species suggests evolutionary constraint, and species-specific molecular evolution of enhancers can help identify gene regulatory sequences that underly the evolution of species-specific traits (*5–8*).

Elements broadly conserved but specifically divergent in humans are called human accelerated regions (HARs) (Figure 1a). As most HARs are noncoding (96%) and 30% of them are predicted to be developmental enhancers, HAR evolution may underlie human-specific gene regulation and attributes including neuroanatomy (*9–12*). Since humans males differ from the males of certain other ape species for traits like sperm production, sperm motility and male aggression (*13–15*), we hypothesized that some HARs underlie evolution of human male-specific traits.

**Figure 1.**
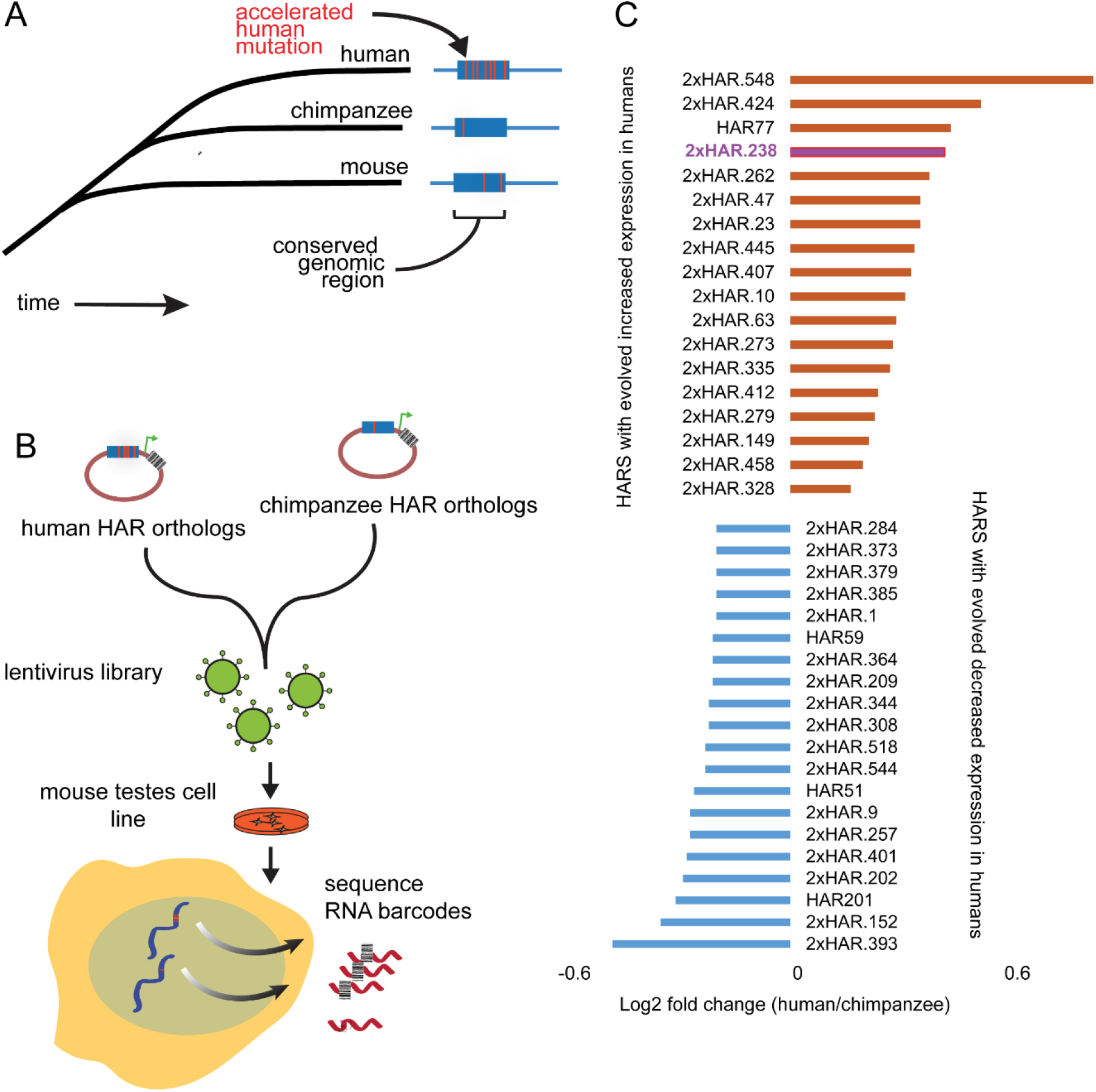
Discovery of putative human-evolved testis enhancers using an MPRA. **(A)** HARs are conserved across mammalian evolution but evolved at an accelerated rate in the human lineage. **(B)** A lentiviral library containing human and chimpanzee HAR orthologs was used to infect GC-1 testis cells for the MPRA. **(C)** HARs identified by the MPRA with an FDR < 0.05 and a fold-change in enhancer activity >0.2 between the human and chimp orthologs are shown. The red bar marks 2xHAR.238.

In addition to the male germline, vertebrate testes include somatic cells that regulate gametogenesis and sexual development through endocrine and paracrine signals. The two principal somatic cells are the Sertoli cells, located in the seminiferous tubules and which form the blood-testis barrier, and the Leydig cells, located in the interstitial compartment and which generate androgens (*16*). We have previously identified a cell line, GC-1, with a transcriptome that is similar to testis somatic cells (*17*).

A key regulator of the function of testis somatic cells is Hedgehog signaling. One Hedgehog, Desert hedgehog (DHH), is produced by Sertoli cells, and signals to Leydig cells (*18–21*). Ablation of mouse *Dhh* disrupts testis development (*19, 21–23*). Interestingly, the effect of loss of DHH on the testis depend on strain, suggesting that genetic modifiers affect testis Hedgehog signaling.

To identify HARs that could be involved in male-specific trait development, we used a high-throughput method to compare enhancer activity of human and chimpanzee HAR orthologs, expressed in the GC-1 testis somatic cell line. We further characterized one HAR, 2xHAR.238, which showed increased activity in the human ortholog relative to the chimpanzee ortholog. 2xHAR.238 is adjacent to *Gli2*, a transcriptional effector of Hedgehog signaling (*12, 22–25*). We found that 2xHAR.238 regulates Leydig cell expression of *Gli2* and physically interacts with the *Gli2* promoter, indicating that it is a *Gli2* enhancer. Deletion of 2xHAR.238 in mice altered inter-male social behavior as well as normal male interest in female odor, implicating this HAR in the evolution of social behavior.

## RESULTS

### A high-throughput screen to identify human-evolved testes gene enhancers

To identify HARs that could regulate male-specific trait development, we used a lentivirus-based massively parallel reporter assay (lentiMPRA) to test the enhancer activity of both the human and chimpanzee orthologs of 714 HARs in a testis somatic cell line (*6, 17, 26–29*). More specifically, sequences containing the 714 HARs were synthesized and cloned, along with a unique barcode, into a lentiMPRA enhancer vector upstream of a minimal promoter driving expression of GFP. Lentivirus libraries were generated and used to infect GC-1 cells. For each human or chimpanzee HAR sequence, we quantified integration via DNA barcode sequencing and determined the relative normalized expression by RNA barcode sequencing followed by calculation of the RNA/DNA log-ratio (Figure 1b; see Methods).

The MPRA identified 18 HARs where the human sequence drove increased expression relative to the chimpanzee ortholog, and 20 which induced decreased expression (Figure 1c). To further test whether the lentiMPRA faithfully identifies regulatory elements with enhancer activity, we assayed the enhancer activity of eight HARs adjacent to genes that regulate testes function (Table 1) using a complementary luciferase assay. Briefly, the sequences of the differentially active HARs were inserted immediately upstream of a firefly luciferase vector promoter. These vectors were co-transfected with a Renilla luciferase vector into GC-1 cells, and enhancer-driven expression was determined from the ratio of firefly to Renilla signal (see Methods). This assay confirmed that three of the tested HARs exhibit different enhancer activity than their chimpanzee orthologs (Figure 2).

**Figure 2.**
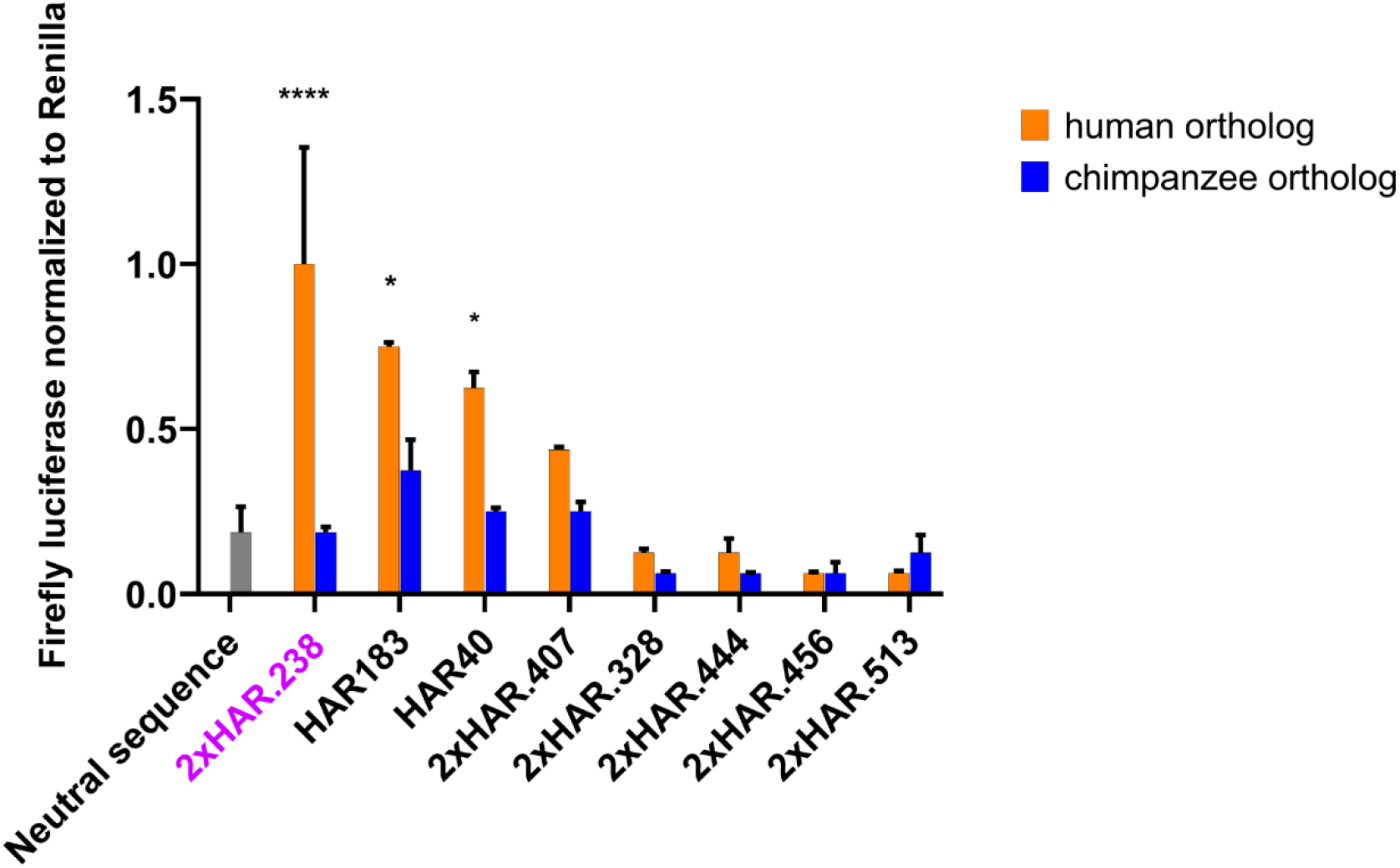
Select HARS and their chimpanzee orthologs identified by MPRA were assayed for their ability to induce luciferase expression in GC-1 cells. Error bars are S.E.M. P-values calculated by Student’s t-test.

**Table 1.**
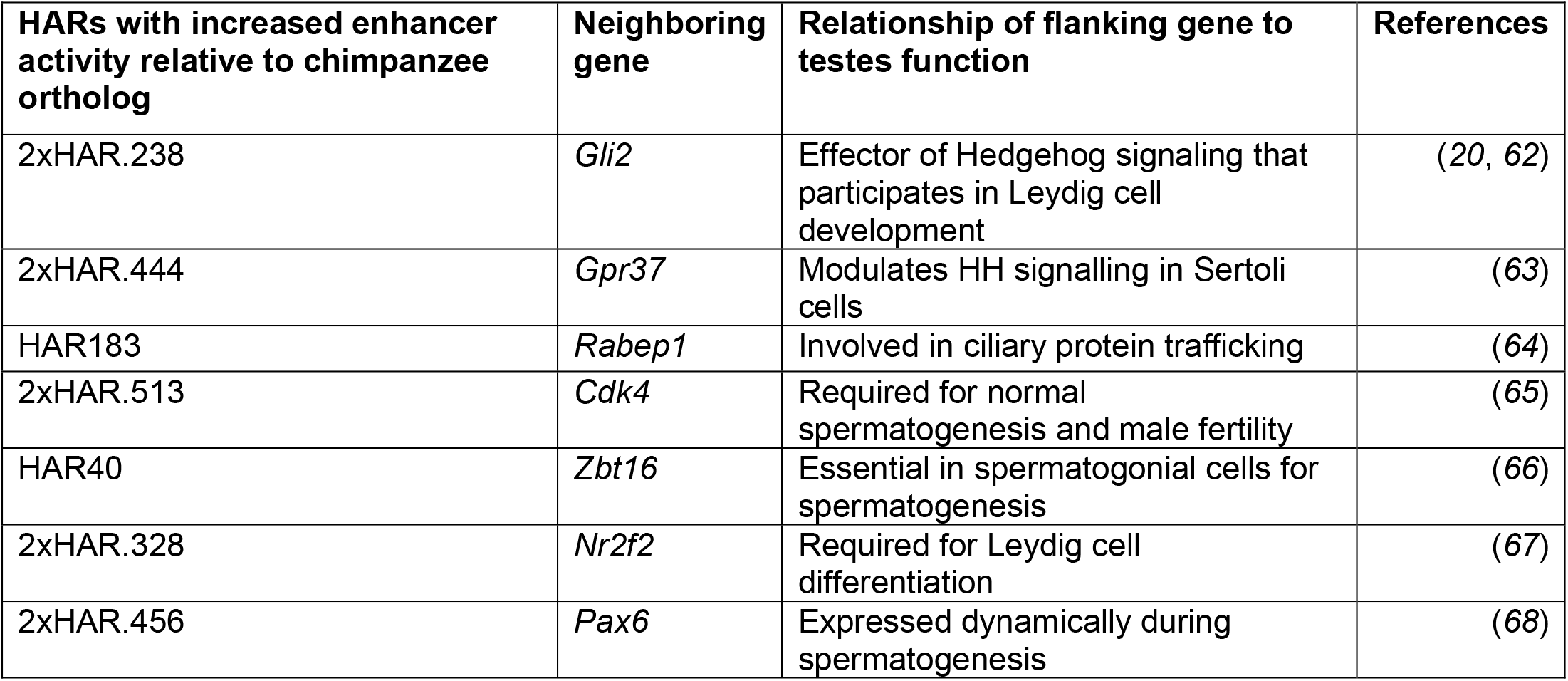

| **HARs with increased enhancer activity relative to chimpanzee ortholog** | **Neighboring gene** | **Relationship of flanking gene to testes function** | **References** |
| --- | --- | --- | --- |
| 2xHAR.238 | *Gli2* | Effector of Hedgehog signaling that participates in Leydig cell development | (20, 62) |
| 2xHAR.444 | *Gpr37* | Modulates HH signalling in Sertoli cells | (63) |
| HAR183 | *Rabep1* | Involved in ciliary protein trafficking | (64) |
| 2xHAR.513 | *Cdk4* | Required for normal spermatogenesis and male fertility | (65) |
| HAR40 | *Zbt16* | Essential in spermatogonial cells for spermatogenesis | (66) |
| 2xHAR.328 | *Nr2f2* | Required for Leydig cell differentiation | (67) |
| 2xHAR.456 | *Pax6* | Expressed dynamically during spermatogenesis | (68) |

### A human-accelerated region regulates *Gli2* expression

One of these HARs, 2xHAR.238, is located 283 kilobases from the transcriptional startsite of *GLI2*, a critical effector of the Hedgehog signaling pathway (*24*). Given the importance of the Hedgehog signaling pathway for male reproductive function (*21–23*), and previous evidence that 2xHAR.238 is a developmental enhancer with human-chimpanzee expression differences (*12*), we assessed the function of 2xHAR.238 in mouse testes.

In many cases, enhancers physically contact the promoters that they influence. To test whether 2xHAR.238 contacts the *Gli2* promoter, we performed chromosome chromatin confirmation capture with sequencing (4C-seq) on mouse GC-1 cells. 4C-seq revealed that 2xHAR.238 interacts with the *Gli2* promoter region (Figure 3a), consistent with a role for 2xHAR.238 as an enhancer of *Gli2*. Interestingly, a previous analysis of human foreskin fibroblasts using Hi-C (*30*) also suggested an interaction between 2xHAR.238 and the *Gli2* promoter (Figure S1). Furthermore, expression of *Gli2* in the developing cortex overlaps 2xHAR.238 expression in transient transgenic mouse reporter assays (*12*). Together, these results suggest that 2xHAR.238 may function as a *Gli2* enhancer.If 2xHAR.238 is an enhancer of *Gli2* in testes somatic cells, then its disruption will reduce *Gli2* expression. Using CRISPR/Cas9, we deleted the 2xHAR.238 locus in GC-1 cells and measured *Gli2* expression (Figure 3b). In 2xHAR.238-deleted cells, *Gli2* expression was reduced, indicating that the 2xHAR.238 sequence is necessary for full *Gli2* expression in GC-1 cells, a testis somatic cell line. Thus, as 2xHAR.238 physically interacts with the *Gli2* promoter and activates *Gli2* expression, we conclude that 2xHAR.238 is an enhancer of *Gli2*.

**Figure 3.**
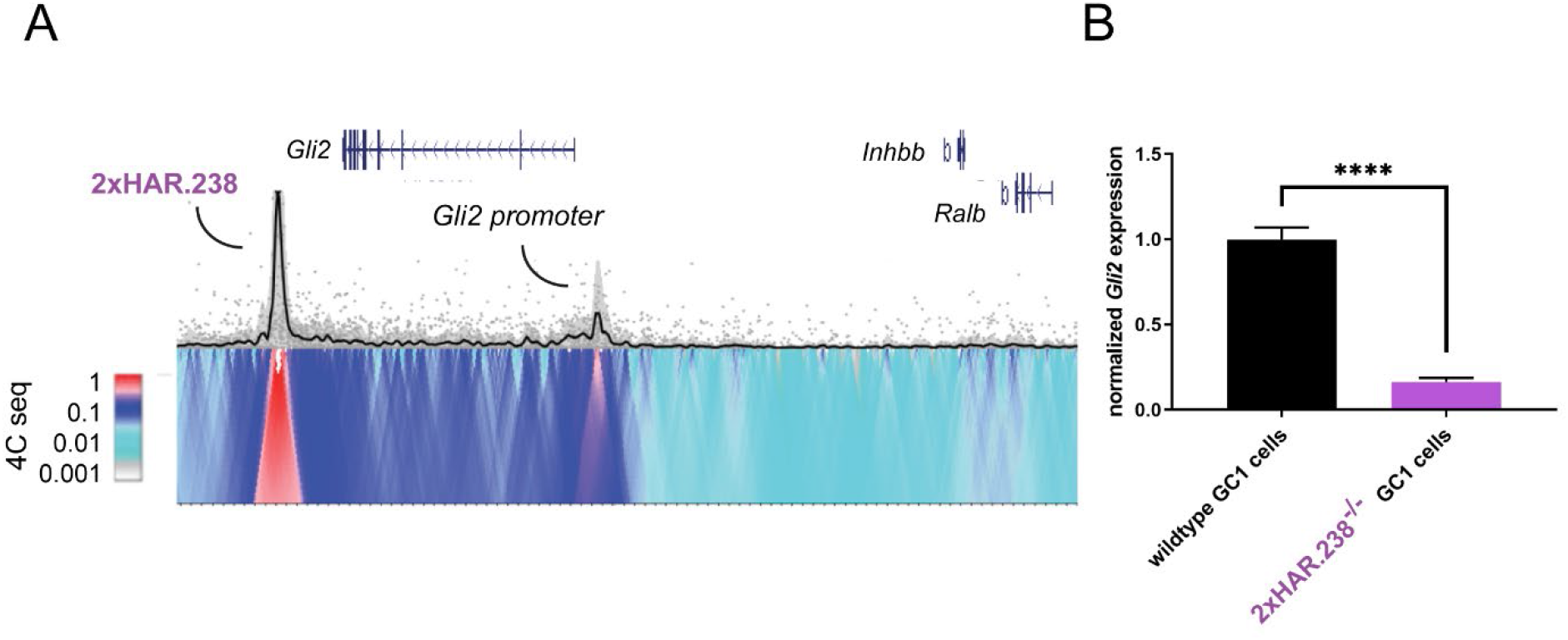
A human accelerated region is a testis enhancer of *Gli2*. **(A)** 4C-seq contact profile of 2xHAR.238 in GC-1 cells. **(B)** qRT-PCR for *Gli2* expression in GC-1 control cells and GC-1 cells in which 2xHAR.238 was deleted. Error bars are S.E.M. P-value from Student’s t-test.

### Mouse 2xHAR.238 regulates *Gli2* expression in neonatal testes Leydig cells

The attenuation of *Gli2* expression upon deletion of 2xHAR.238 in the GC-1 cells suggested that the enhancer might regulate expression in the testes. To test this hypothesis, we deleted 2xHAR.238 using CRISPR/Cas9 in mice by co-injection of Cas9 and guide RNAs targeted to sequences flanking the HAR (see Methods). 2xHAR.238^-/-^ mice were viable and exhibited no gross defects in body or testes morphology.

As GC-1 cells are testes somatic cells, similar to immature Leydig cells, and Leydig cells require Hedgehog signaling for maturation (*17, 21*), we investigated the possibility that the 2xHAR.238 enhancer regulates the expression of *Gli2* in Leydig cells. Using in situ hybridization, we assayed the expression of *Gli2* in the testes of 2xHAR.238^-/-^ and wild type newborn mice. We simultaneously examined the expression of Cyp17A1, a marker of Leydig cells. *Gli2* expression was reduced specifically in the Leydig cells of 2xHAR.238^-/-^ perinatal testes (Figure 4a,b; quantification in Figure 4c), indicating that 2xHAR.238 is required for normal *Gli2* expression in Leydig cells in vivo. To test the ability of 2xHAR.238 to drive expression of a transgene in testes, we generated mice carrying 2xHAR.238 adjacent to a minimal promoter and upstream of *lacZ*. We assayed testes of newborn mice for *lacZ* transgene by in situ and found expression specifically in the testis somatic cells, and not the germ cells (Figure 4d,e; quantification in Figure 4f). Together, these results indicate that the 2xHAR.238 enhancer is necessary for normal *Gli2* expression in Leydig cells, and sufficient to drive transgene expression in testis somatic cells.

**Figure 4.**
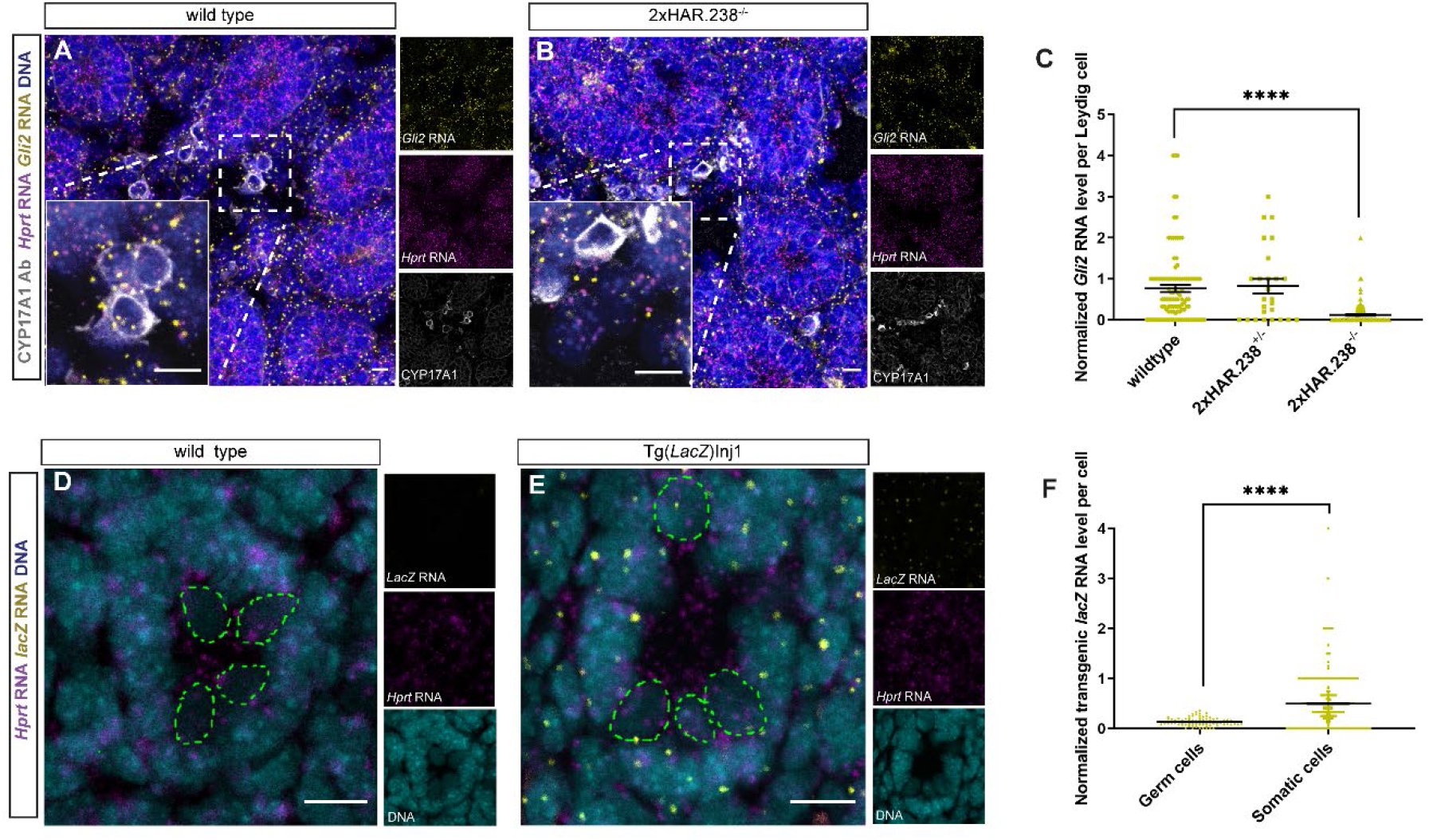
2xHAR.238 regulates expression of *Gli2* in newborn mouse Leydig cells. (A) Wild type and (B) 2xHAR.238^-/-^ newborn (P6) mouse testes sections assessed for *Gli2* expression (yellow) and control message (*Hprt*, magenta) by RNAscope in situ hybridization. CYP17A1 immunofluorescence (white) indicates Leydig cells. Hoechst staining indicates nuclei (blue). Insets are magnified views. (C) Quantitation of *Gli2* expression normalized to *Hprt* expression in Leydig cells. n = 2 wild type, 1 2xHAR.238^-/+^ and 2 2xHAR.238^-/-^ mice. (D) Wildtype and (E) Tg(*lacZ*)Inj1 transgenic newborn mouse (P6) testes sections assessed for *lacZ* expression (yellow) and control message (*Hprt*, magenta) by RNAscope. Cyan is Hoechst. (F) Quantification of *lacZ* expression normalized to *Hprt* expression in the germ and somatic cells of a single mouse testis. Germ cells are the round, lightly staining nuclei in the testis tubule lumen, denoted by the dotted green line. Error bars are S.E.M. P-values calculated by Welch’s t-test.

### 2xHAR.238 function is involved in male-typical behaviors

In mice, perinatal Leydig cell function enable male-typical behaviors such as home-cage aggression (*31, 32*). To investigate the possible function of 2xHAR.238 in inter-male aggression, we employed a standard rodent resident intruder assay (see Methods). Briefly, 2xHAR.238^-/-^ mice and wild type controls (“residents”) were placed in a cage for one day to establish a home territory.

On days 2, 3 and 4, the residents were challenged with an intruder male and responses were recorded. These data were expressed as an intruder dominance score, which is a measure of the total bouts of following, sniffing, mounting, licking, circling, tail grabs and pushing during a 10-minute period (Figure 5a). Interestingly, on day 4, 2xHAR.238^-/-^ males exhibited increased passivity in response to male intrusion into their home cage (Figure 5b).

**Figure 5.**
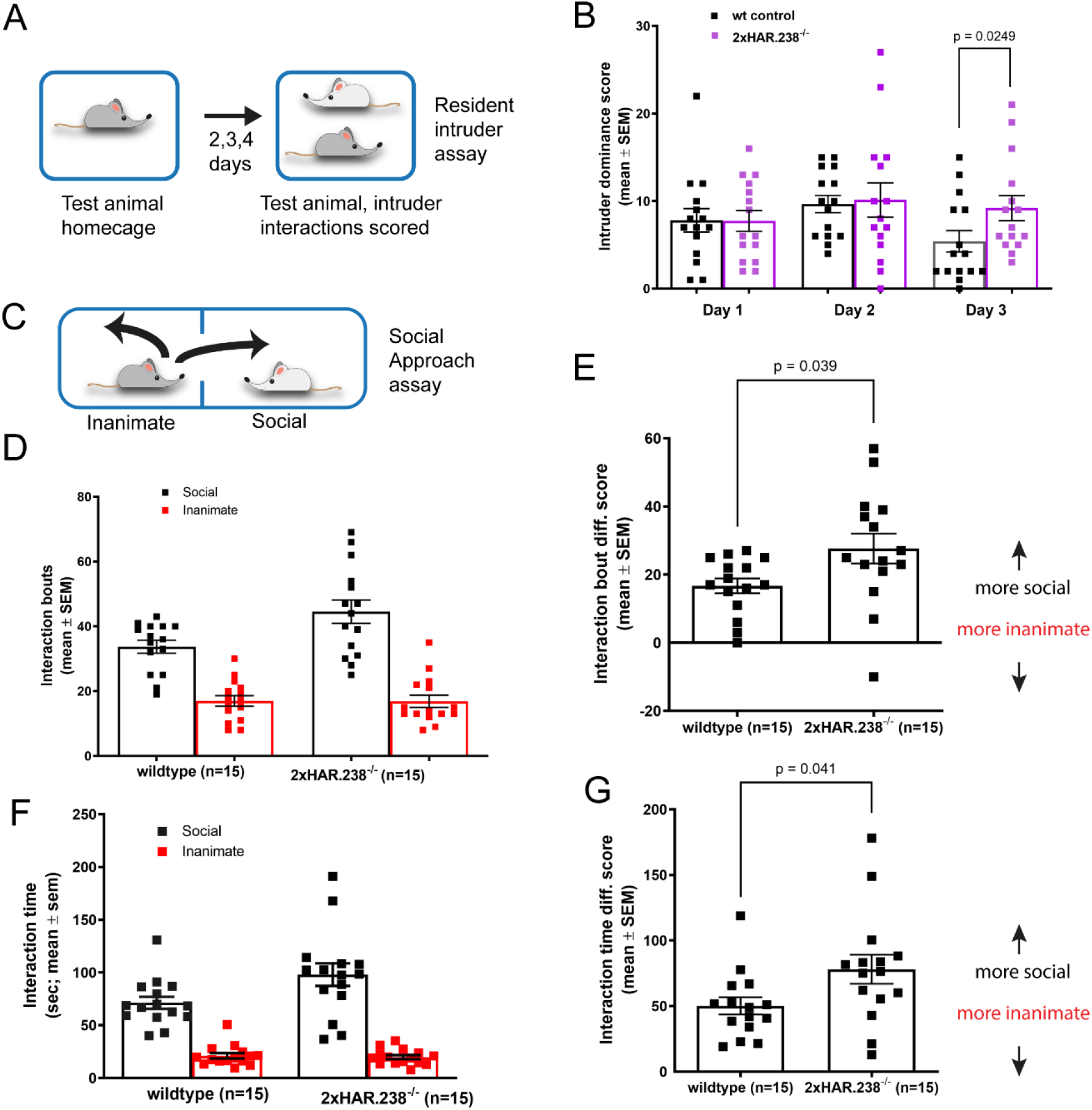
2xHAR.238 is involved in inter-male social behavior. (A) In the resident intruder assay, the experimental subject mouse establishes a home cage and is challenged by an intruder male on three subsequent days. Mounting, sniffing, licking, circling, tail grabs, and pushing by the intruding male are scored. (B) On day 3, male mice intruding on *2xHAR.238^-/-^* mice engaged in more of these dominance behaviors than those intruding on wild type control males. P-value for the resident intruder assay was calculated using the Mann Whitney test. (C) In the social approach assay, the experimental subject mouse remains in the “inanimate” chamber or moves to the “social chamber”, where a stimulus mouse is confined. (D,E) *2xHAR.238^-/-^* mice engaged in more interaction bouts with the stimulus mouse than did wild type control mice. “Interaction bout difference score” is the difference between the interaction bouts in the two chambers for the given genotype. (F,G) *2xHAR.238^-/-^* mice also spent more time interacting with the stimulus mouse. “Interaction time difference score” is the difference between the interaction times in the two chambers for the given genotype. P-values for the social approach assay were calculated using Welch’s t-test.

To further assess the inter-male behavior of 2xHAR.238^-/-^ males, experimental mice were tested using the standard social approach assay. In this assay, a second male mouse is placed in a cage adjoining that of the subject male mouse with free airflow, and the subject mouse can choose to investigate the novel male mouse or retreat (Figure 5c). Consistent with altered intermale behavior, 2xHAR.238^-/-^ mice were more likely than wild type controls to socially interact with the novel male, both in terms of number of interaction bouts and the mean interaction time per bout (Figure 5d-g). Importantly, subject mice did not display altered locomotion or olfaction (Figure S2a,b).

In addition to control of inter-male behavior, Leydig cell function is important for male-typical mating behavior (*32*). To assess the interest of male mice in female odor, male 2xHAR.238^-/-^ or wild type mice were presented with water, male cage odor or female cage odor during an odor habituation test (see Methods). Interestingly, while there was no significant difference between the wildtype control males and 2xHAR.238^-/-^ males when presented with water or male odor, 2xHAR.238^-/-^ males interacted with the female odor for less time (Figure 6a) and for fewer bouts (Figure 6b) than did control males.

**Figure 6.**
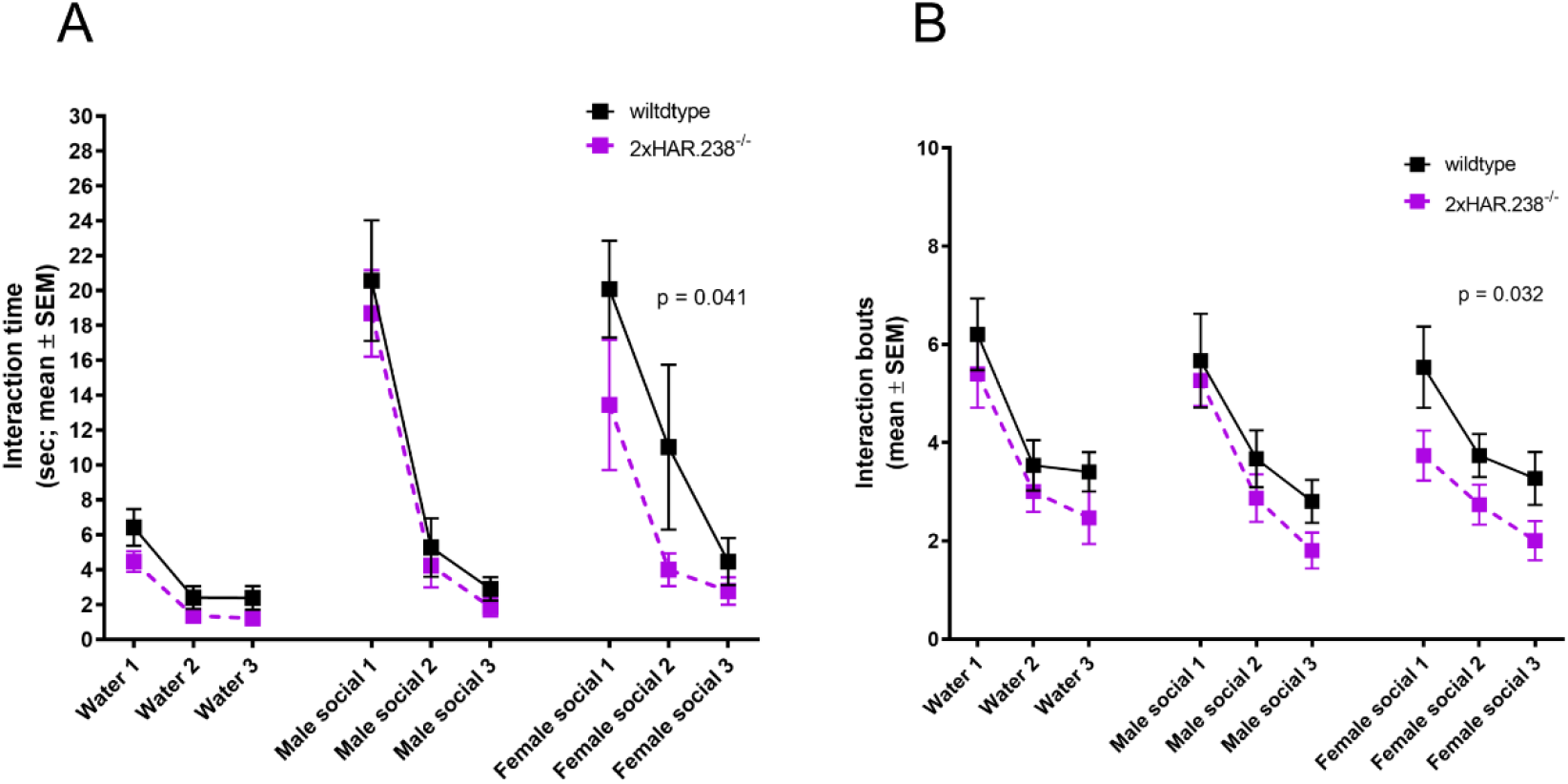
2xHAR.238^-/-^ male mice habituate more quickly to female odor. Both 2xHAR.238^-/-^ and wild type control males distinguish between water and mouse bedding odor (social odor). However, 2xHAR.238^-/-^ male mice displayed faster habituation to female social odor than did wild type control male mice, both in terms of time interacting with female odor (A) and number of interaction bouts with presented odor per trial (B). n = 15 for both genotypes.

## DISCUSSION

HARs are genomic regions conserved through mammalian evolution, but which have undergone accelerated evolution, specifically in the human lineage. Using an MPRA, we identified a set of HARs that are testis cell enhancers. One of these enhancers, 2xHAR.238, controls the expression of the Hedgehog pathway effector *Gli2*, both in a testes somatic cell line and in perinatal testis Leydig cells. 4C-seq revealed that 2xHAR.238 interacts with the *Gli2* promoter region. Deletion of 2xHAR.238 disrupted normal mouse inter-male behavior. As 2xHAR.238 is both a HAR and an enhancer of *Gli2*, a cis-regulatory element of *Gli2* has evolved uniquely in the human lineage. Therefore, we hypothesize that evolution of regulatory regions controlling testes gene expression may have affected the evolution of human behavior.

Deletion of 2xHAR.238 reduced *Gli2* expression in the Leydig cells of the testes during the perinatal period, indicating that this enhancer promotes *Gli2* expression. 2xHAR.238 also drove expression of a transgene in the newborn testis somatic cells, consistent with its in vitro enhancer activity. During perinatal testis development, Desert hedgehog is required for the maturation of Leydig cells (*21, 22*). As the main transcriptional effector of Hedgehog signaling, GLI2 is likely to interpret Desert hedgehog signals to the Leydig cells.

In the immediate post-natal period of males, testosterone is synthesized by fetal Leydig cells to generate an early surge of androgens, sometimes referred to as a “mini-puberty”, which is required for the development of male-specific behavior (*33–36*). Consequently, conditional inactivation of the androgen receptor in the mouse nervous system diminishes some forms of masculine behavior, such as aggression (*37*). We found that deletion of 2xHAR.238 compromises some forms of inter-male behavior in adult mice, including decreasing inter-male aggression and increasing inter-male sociality. Therefore, 2xHAR.238-dependent regulation of *Gli2* expression may contribute to adult male-specific behavior via control of androgen synthesis. Further investigation, using tissue-specific inactivation of *Gli2* and perinatal androgen measurements, will be necessary to test this hypothesis.

2xHAR.238 is an enhancer that affects mouse behavior. A classic example of an indispensable enhancer is the ZRS, which regulates the Hedgehog pathway via *Sonic hedgehog* (*38*). Loss of ZRS function disrupts limb development in mice and underlies the evolution of reduced limbs in snakes (*3*). Furthermore, enhancer evolution leading to decreased expression of the Hedgehog receptor *Ptch1* may underly the evolution of bovine limbs (*39*). Alteration in genes encoding components of primary cilia, involved in transducing Hedgehog signals, may affect wing shape in flightless Galapagos cormorants (*40*). Our results indicate that another member of the Hedgehog pathway, *Gli2*, is regulated by a critical, evolved enhancer (Figure 7). Together, these findings reveal the Hedgehog pathway to be a frequent target of species-specific Chimpanzee males engage in more non-lethal aggression than hunter-gatherer human males (*15*), suggesting that male-specific behavior has evolved in the human lineage relative to other great apes. The molecular evolution of a HAR, 2xHAR.238, could be related to this change in human behavior. Further investigation, including analysis of mice humanized at the 2xHAR.238 locus, will be necessary to test these possibilities. In addition, other HARs identified in this study may illuminate other aspects of human male-specific gene regulation. For example, another HAR with increased differential activity in humans and chimpanzees, 2xHAR.328, adjoins *NR2F2*, better known as *COUP-TFII*, which cooperates with GLI2 to regulate Leydig cell androgen production. Thus, multiple HARs may underlie the evolution of human male-specific biology.

**Figure 7.**
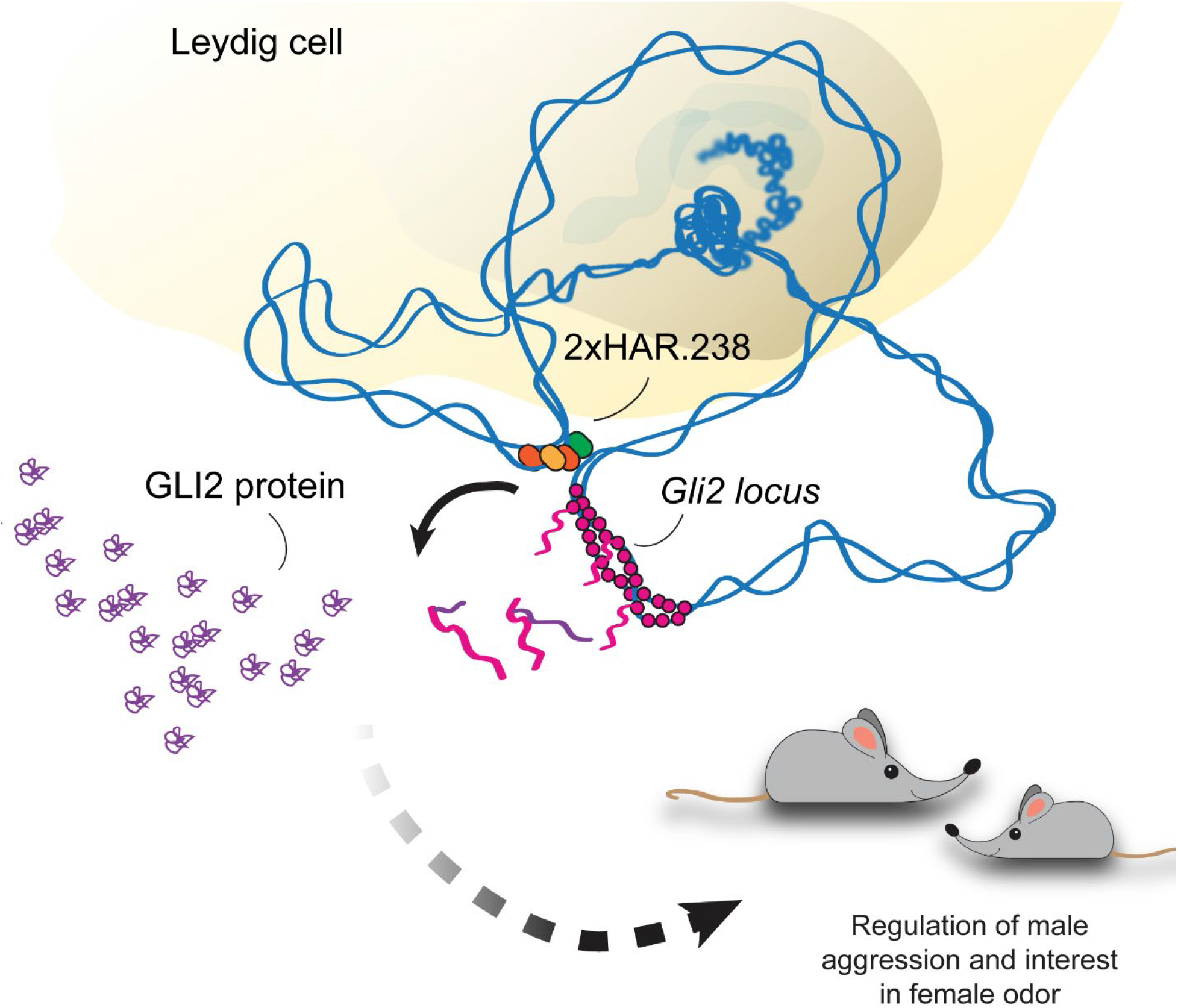
A human accelerated region is an enhancer of *Gli2* in Leydig cells and affects male social behavior.

## METHODS

### MPRA

Library construction, infection and sequencing: Library design/synthesis/cloning and lentivirus library packaging was carried out as described in (*9*). Infection with lentiviral packaged library and DNA/RNA isolation from GC-1 cells and sequencing was conducted in triplicate, and carried out as described in (*9*).

#### Analysis

RNA and DNA reads were mapped and counted, so that expression of each construct in the library could be normalized to genomic integration. The ratio of RNA reads to DNA reads (ratio score) for each HAR construct was calculated individually for each of the three replicates. Since each HAR construct was represented by multiple fragments, the fragment from each HAR construct with the maximum ratio score was selected to represent the ratio score for that HAR construct. This was done separately in each library replicate, across both human and chimp HAR construct orthologs. Limma (*41*) linear modeling was used to determine, for each HAR, if the ratio scores observed in one species ortholog across replicates was significantly different from the ratio score observed for the other species. P-values were adjusted for multiple testing using the Benjamini-Hochberg false discovery rate (FDR) correction. This value was reported as the negative log of the FDR estimate (*42*).

### Luciferase Assay

Significant results from the MPRA assay of interest were amplified by PCR and cloned into the pLS-mP-Luc vector (Addgene Cat# 106253) in place of GFP. These included the human and chimpanzee orthologs of 2xHAR.513 (hg19 chr12:59202762-59202994), 2xHAR.238 (hg19 chr2:121549404-121549483), 2xHAR.328 (hg19 chr15:94084865-94085048), 2xHAR.407 (hg19 chr3:140176936-140176987), 2xHAR.444 (hg19 chr7:124454421-124454447), 2xHAR.456 (hg19 chr11:31993804-31993909), HAR183 (hg19 chr17:6050354-6050576), HAR40 (hg19 chr11:114099775-114100193) and a neutral sequence (hg19 chr2:124107843-124108061) showing no evidence of enhancer activity based on publicly available ENCODE ChIP data (*43*). The renilla control vector used was from (*9*). Lentivirus was generated using 10:1 molar mixtures of firefly-luciferase experimental to renilla-luciferase control vectors for each HAR listed above using a standard protocol outlined in (*44*). GC-1 cells were seeded in a well at 1/10 density and a well of cells was infected with a lentivirus mixture for each tested HAR, in triplicate, 24 hours after seeding. Lucerifase measurements were taken on a Promega Glomax plate reader after sample preparation using the Promega Duel-Luciferase Reporter System (Promega Cat# E1910) 72 hours after lentivirus infection. Enhancer activity was calculated as the ratio of firefly to renilla activity for each construct.

### Enhancer deletions

#### GC-1 cells

Crispr/Cas9 expression constructs were made using the PX459 plasmid (Addgene Cat# 62988) as described in (*45*). Crispr small guide RNAs (sgRNAs) were selected using the GPP sgRNA Designer (*46*). A construct targeting the left flank of 2xHAR.238 was made using the sgRNA sequence GCTGCAGAGAGTCCATTAGA (mm10) and another targeting the right flank using the sequence TGTGATTAATACACTTGCCC (mm10). The sgRNA sequences were selected to maximize predicted on-target cutting and minimize predicted off-target cutting. These two constructs were co-transfected in GC-1 mouse testes cells at 80% confluency with Lipofectamine 3000 (Invitrogen Cat# L3000001) using 1.5μL of Lipofectamine and 250 ng of each vector in a well of a 24-well plate. After 2 days these cells were passaged and treated with puromycin (3μg/mL, Thermofisher Cat# A1113802) for an additional 2 days to reduce the number of un-transfected cells in the culture. Cells in the culture were counted and passaged onto 10 cm cell culture plates at a density of 1000 cells per plate. After 2 weeks, colonies were picked, expanded and Sanger sequenced for genotype using amplicons from PCR primers TAAGCCTAGCATTACAAGAAGGTG and CAATACTCAGACTGTTGGTGGAAAC.

#### Mice

All animal procedures were approved by the Institutional Animal Care and Use Committee at the University of California, San Francisco. Super-ovulated female C57BL/6 mice (4 weeks old) were mated to C57BL/6 stud males. Fertilized zygotes were collected from oviducts. Cas9 protein (30 - 60 ng/ul) and the same 2xHAR.238-flanking sgRNA species (15-30 ng/ul) used in the GC-1 deletion experiment were co-injected into the pronucleus of fertilized zygotes. Injected zygotes were implanted into oviducts of pseudopregnant CD1 female mice. Pups were genotyped by Sanger sequencing in the same manner as the GC-1 cells. The 2xHAR.238-deleted mouse line was maintained through intercross of 2xHAR.238^-/-^ animals.

### Transgenic Enhancer Assay

The mouse ortholog of 2xHAR.238 was inserted upstream of the hsp68 minimal promoter in a lacZ expression plasmid and transgenic mice were generated as described in (*47*).

### qRT-PCR

RNA was isolated from cells using the Qiagen RNeasy Mini Kit (Cat# 74104) and cDNA was prepared using the iScript cDNA Synthesis Kit (Biorad, Cat# 1708890). Assays were done in triplicate using Express SYBR GreenER master mix (Invitrogen) and an ABI 7900HT RT-PCR machine (Applied Biosystems). Expression levels were normalized to the expression level of the housekeeping gene *HMBS* and quantitative changes were calculated using the method described in (*48*). The primers used for *Gli2* were CCGAGGTGGTCATCTACGAG and AACTCCTTCTTCTCCCCGTG. The primers used for *HMBS* were CAGAGAAAGTTCCCCCACCT and GGATGATGGCACTGAACTCC. qRT-PCR primers were designed using the qPCR Primers track on the UCSC genome browser (*49*).

### 4C-seq

2×10^7^ cells were fixed in PBS with 2% formaldehyde for 10 minutes at RT. Crosslinking was quenched with 125 mM Glycine. The cells were centrifuged and resuspended in lysis buffer (10 mM Tris-HCl pH7.5, 10 mM NaCl, 5 mM MgCl2, 100 mM EGTA, 1x complete proteinase inhibitors (Roche, Cat# 11697498001)) and incubated for 5 minutes on a nutator. Cells were then centrifuged and washed with lysis buffer. The cell pellet was stored at −80°C until use. The cell pellet was suspended in 0.5% SDS and incubated at 62°C for 10 min. 10% Triton X-100 was added to the pellet and incubated at 37°C for 15 min. Chromatin was digested with 100 units of *Nla*III (New England Biolabs, Cat# R0125) at 37°C overnight. After heat inactivation of the enzyme, 2,000 units of T4 DNA ligase (New England Biolabs, Cat# M0202) was added and incubated at room temperature for 4 hours. The ligated chromatin was pelleted and resuspended in EB buffer (Qiagen, Cat# 19086). Proteinase K (New England Biolabs, Cat# P8107) and 10% SDS were added, mixed by inversion and incubated at 55°C for 30 min. 5M NaCl was added and the sample was incubated at 68°C overnight. DNA was purified by SPRISelect beads (B23319) and resuspended in 50 ul EB buffer. DNA was digested with 50 units of *Dpn*III (New England Biolabs, Cat# R0543) at 37°C for 4 hours. Following heat inactivation of the enzyme at 65°C, DNA was purified by SPRISelect beads. A second round of ligation was performed with 400 units of T4 DNA ligase in a 14 ml reaction volume at 16°C overnight. DNA purified using phenol-chloroform and ethanol precipitation. Inverse PCR was performed using NEBNext high-fidelity 2X PCR master mix (New England Biolabs, catalog no. M0541). DNA was purified with SPRISelect beads. Massively parallel sequencing was performed on an Illumina HiSeq4000 with 50 bp single-end reads or Illumina NovaSeq with 150 bp paired-end reads that were subsequently trimmed to 50 bp for data analysis. 4C-seq was carried out using two biological replicates. 4C-seq data was analyzed using the 4C-seq pipeline (*50*). The primers used for the PCR were:

FP for HAR (4C): AATGATACGGCGACCACCGAGATCTACACTCTTTCCCTACACGACGCTCTTCCGATCTGTGTGATGCCCAAGGGCATG
RP for HAR (4C): CAAGCAGAAGACGGCATACGAGATTCCGTACGGTGACTGGAGTTCAGACGTGTGCTCTTCCGATCTAAATCACGGCAGATGGCAGG

### In situ hybridization

Testes were harvested from mouse pups from P0 to P6 and fixed in 4% paraformaldehyde in PBS overnight at 4°C. Testes were washed in PBS, incubated in 30% sucrose in PBS overnight at 4°C and embedded in OCT (Tissue-Tek). Twelve micron sections were cut from the embedded blocks and used for subsequent in situ hybridizations with the RNAscope Multiplex Fluorescent Reagent Kit v2 (ACD, Cat# 323110), as described in (*51*) with the following modifications: slides were washed twice with PBS at the start of the protocol and dehydrated in 100% ethanol for 3 minutes; antigen retrieval was performed for 15 minutes; RNAscope Protease IV was applied to the sections for less than 1 minute at RT, and immediately washed twice in PBS before beginning the hybridization procedure. The in situ probes used were Mm-Gli2 (ACD, Cat# 405771), Mm-Hprt-C2 (ACD, Cat# 312951-C2), and Ecoli-LacZ (ACD, Cat# 461191). For sections co-stained with antibody, slides were blocked (2% BSA/1% donkey serum in PBS) for 1 hour at RT immediately following RNA scope protocol. Rabbit anti-Cyp17a1 (Proteintech, Cat# 14447-1-AP) was applied at dilution 1:500 for 1 hour at room temperature, followed by application of donkey anti-rabbit 647 secondary antibody (Thermofisher, cat# A-31573) at dilution 1:500 for 45 minutes at RT. Analysis of RNAscope expression level was done by counting RNAscope in situ signal puncta using the Find Maxima function in ImagJ (*52*): channels with probe signal were subjected to Gaussian Blur using ImageJ (radius = 1), puncta identified with Find Maxima (prominence = 20) and maxima within marker-defined Leydig cells counted.

### Behavioral tests

Behavioral assays were conducted by the UCSF Gladstone Mouse Phenotyping Core Facility. All observers were blinded to mouse genotype. C57BL/6 used as experimental animals were aged-matched and bred from 2xHAR.238^-/-^ x 2xHAR.238^-/-^ and 2xHAR.238^+/+^ x 2xHAR.238^+/+^ parents.

#### Open-Field Test

Mice were placed in a clear chamber and habituated for 1 hour, then allowed to explore for 15 minutes while movement was automatically monitored. These data were used to establish a baseline of general locomotor activity, exploration, and anxiety levels. Procedure conducted as described in (*53–55*).

#### Olfactory Habituation Test

Mice were placed in a fresh cage and habituated for 30 minutes. Odors (water, then odor from a male cage, then odor from a female cage) were presented to each mouse on a cotton tip applicator. Time that each mouse spent sniffing each applicator was measured and number of separate investigations with each mouse was counted. Trials were done in triplicate. These data can be used to assess ability to discriminate odors and infer interest in certain types of odors. Procedure conducted as described in (*56, 57*).

#### Two-Trial Social Approach Assay

Experimental mice were placed in a two-chambered enclosure for 1 hour prior to trial for acclimation. An opening allowed free movement between chambers. A male BALB/c stimulus mouse was placed in a sub-enclosure in one chamber (“social”). During the trials, the number of bouts of investigation and time spent investigating the social or non-social chambers was recorded. Greater or less interest in the social chamber can indicate social abnormalities. Procedure conducted as described in (*56, 58, 59*).

#### Resident Intruder Assay

This assay is conducted over 3 days. The experimental mouse (“resident”) is placed in a home cage that is unchanged for one week. On the first day of the assay, a male BALB/c mouse (“intruder”) was placed in the resident home cage for 10 minutes. The occurrence, severity and length of each attack by the resident on the intruder was recorded and scored. Any additional aggressive, defensive and/or fearful behaviors by the resident or the intruder was scored. Note that any excessive attack lasting longer than 30 seconds terminated the trial. On days 2 and 3, the trial was repeated with different intruder mice. The intruder dominance score is the total bouts of following, sniffing, mounting, licking, circling, tail grabs, and pushing during each 10-minute Resident Intruder session. Detail of procedure in (*57, 60, 61*).

## ACKNOWLEDGEMENTS

We thank E Yu for help with animal husbandry and genotyping of mice used in this study. We thank the Gladstone Behavioral Core, especially Jeff Simms and Iris Lo, for their valuable work and expertise. We thank Junli Zhang at the Gladstone ES Cell Targeting Core and the Black Lab at the UCSF Cardiovascular Research Institute for their work and expertise with mouse oocyte injections. We thank members of the Reiter and Ahituv labs for helpful discussions. A.R.N. thanks Brian Black and Pao-Tien Chuang for their helpful insights as members of his thesis committee. This work was supported by NIH R01HD089918 and R01AR054396 to J.F.R., and 1R01MH109907 to N.A. and K.S.P., the Roche/ARCS Foundation Fellowship to A.R.N., and the Gladstone Institutes to K.S.P. J.F.R. and K.S.P. are Chan Zuckerberg Biohub Investigators.

---

**Figure S1.**
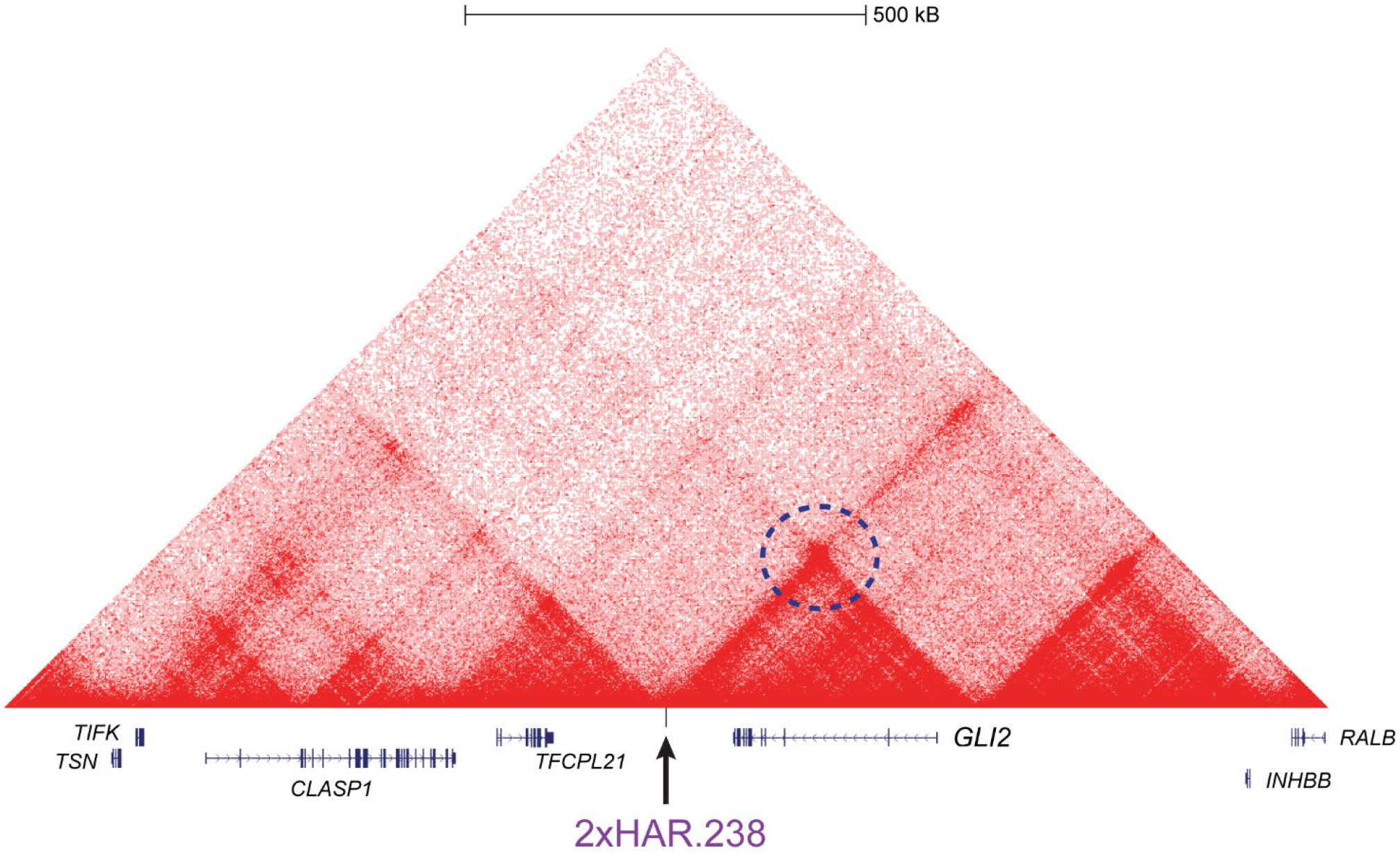
Hi-C analysis of the *GLI2* genomic region in human fetal foreskin fibroblasts. 2xHAR.238 interacts with the promoter region of *GLI2*, marked by the dashed circle. Data are adapted from Krietenstein et al., 2020 (*26*).

**Figure S2.**
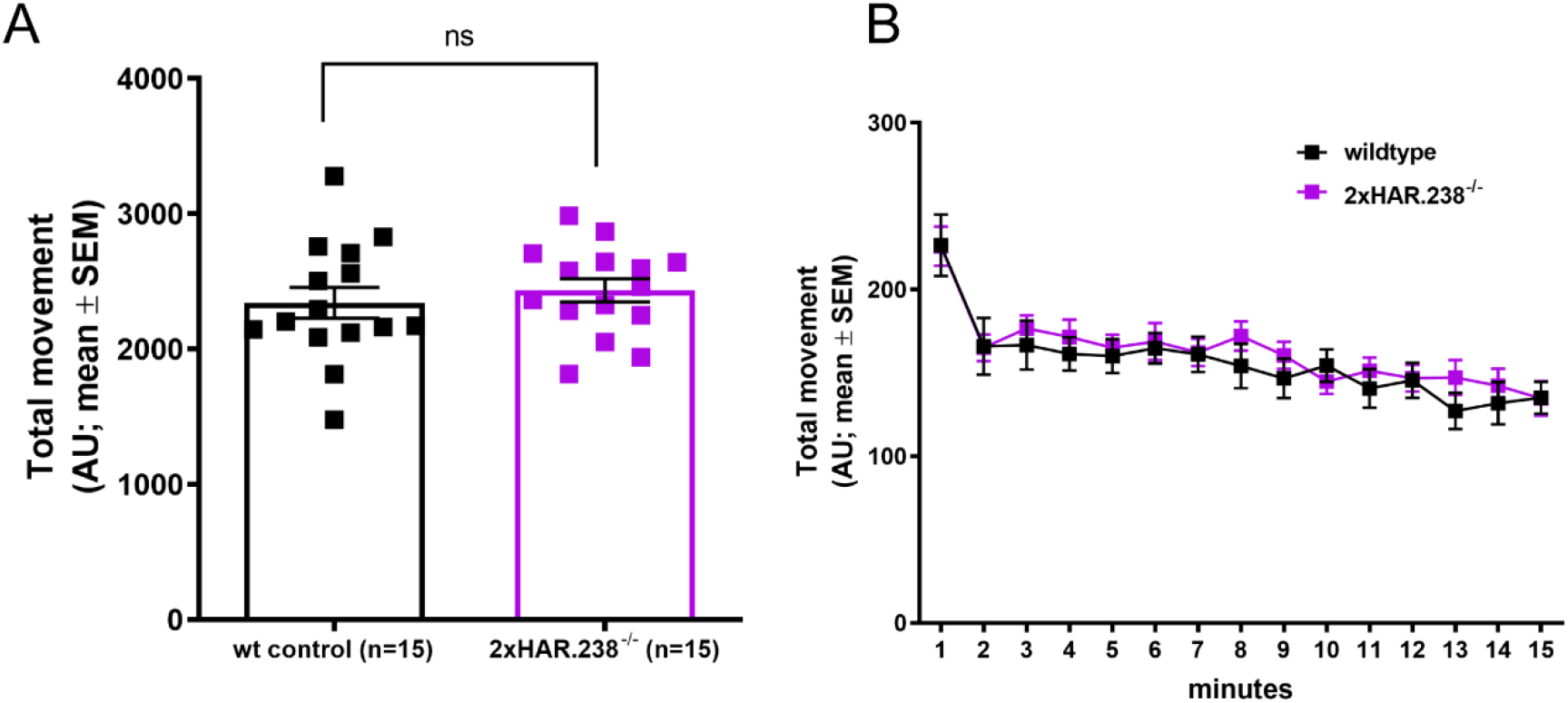
2xHAR.238 deletion does not affect locomotion. (A,B) No difference was observed between *2xHAR.238^-/-^* male mice and wild type controls in the open field activity test, which assays locomotion.

## REFERENCES

1. J.-W. Hong, D. A. Hendrix, M. S. Levine, Shadow enhancers as a source of evolutionary novelty. Science. 321, 1314 (2008).

2. M. Osterwalder, I. Barozzi, V. Tissières, Y. Fukuda-Yuzawa, B. J. Mannion, S. Y. Afzal, E. A. Lee, Y. Zhu, I. Plajzer-Frick, C. S. Pickle, M. Kato, T. H. Garvin, Q. T. Pham, A. N. Harrington, J. A. Akiyama, V. Afzal, J. Lopez-Rios, D. E. Dickel, A. Visel, L. A. Pennacchio, Enhancer redundancy provides phenotypic robustness in mammalian development. Nature. 554, 239–243 (2018).

3. E. Z. Kvon, O. K. Kamneva, U. S. Melo, I. Barozzi, M. Osterwalder, B. J. Mannion, V. Tissières, C. S. Pickle, I. Plajzer-Frick, E. A. Lee, M. Kato, T. H. Garvin, J. A. Akiyama, V. Afzal, J. Lopez-Rios, E. M. Rubin, D. E. Dickel, L. A. Pennacchio, A. Visel, Progressive Loss of Function in a Limb Enhancer during Snake Evolution. Cell. 167, 633–642.e11 (2016).

4. M. Rebeiz, J. E. Pool, V. A. Kassner, C. F. Aquadro, S. B. Carroll, Stepwise Modification of a Modular Enhancer Underlies Adaptation in a Drosophila Population. Science. 326, 1663–1667 (2009).

5. C. Y. McLean, P. L. Reno, A. A. Pollen, A. I. Bassan, T. D. Capellini, C. Guenther, V. B. Indjeian, X. Lim, D. B. Menke, B. T. Schaar, A. M. Wenger, G. Bejerano, D. M. Kingsley, Human-specific loss of regulatory DNA and the evolution of human-specific traits. Nature. 471, 216–219 (2011).

6. K. S. Pollard, S. R. Salama, B. King, A. D. Kern, T. Dreszer, S. Katzman, A. Siepel, J. S. Pedersen, G. Bejerano, R. Baertsch, K. R. Rosenbloom, J. Kent, D. Haussler, Forces shaping the fastest evolving regions in the human genome. PLoS Genet. 2, e168 (2006).

7. C. P. Bird, B. E. Stranger, M. Liu, D. J. Thomas, C. E. Ingle, C. Beazley, W. Miller, M. E. Hurles, E. T. Dermitzakis, Fast-evolving noncoding sequences in the human genome. Genome Biol. 8, R118 (2007).

8. S. Prabhakar, J. P. Noonan, S. Pääbo, E. M. Rubin, Accelerated evolution of conserved noncoding sequences in humans. Science. 314, 786 (2006).

9. H. Ryu, F. Inoue, S. Whalen, A. Williams, M. Kircher, B. Martin, B. Alvarado, Md. A. H. Samee, K. Keough, S. Thomas, A. Kriegstein, J. Shendure, A. Pollen, N. Ahituv, K. S. Pollard, “Massively parallel dissection of human accelerated regions in human and chimpanzee neural progenitors” (preprint, Evolutionary Biology, 2018), doi:10.1101/256313.

10. R. N. Doan, B.-I. Bae, B. Cubelos, C. Chang, A. A. Hossain, S. Al-Saad, N. M. Mukaddes, O. Oner, M. Al-Saffar, S. Balkhy, G. G. Gascon, Homozygosity Mapping Consortium for Autism, M. Nieto, C. A. Walsh, Mutations in Human Accelerated Regions Disrupt Cognition and Social Behavior. Cell. 167, 341–354.e12 (2016).

11. A. Fuentes, How Humans and Apes Are Different, and Why It Matters. Journal of Anthropological Research. 74, 151–167 (2018).

12. J. A. Capra, G. D. Erwin, G. McKinsey, J. L. R. Rubenstein, K. S. Pollard, Many human accelerated regions are developmental enhancers. Philos. Trans. R. Soc. Lond., B, Biol. Sci. 368, 20130025 (2013).

13. M. J. Anderson, A. F. Dixson, Sperm competition: motility and the midpiece in primates. Nature. 416, 496 (2002).

14. H. Fujii-Hanamoto, K. Matsubayashi, M. Nakano, H. Kusunoki, T. Enomoto, A comparative study on testicular microstructure and relative sperm production in gorillas, chimpanzees, and orangutans. Am. J. Primatol. 73, 570–577 (2011).

15. R. W. Wrangham, M. L. Wilson, M. N. Muller, Comparative rates of violence in chimpanzees and humans. Primates. 47, 14–26 (2006).

16. S. Schlatt, J. Ehmcke, Regulation of spermatogenesis: an evolutionary biologist’s perspective. Semin Cell Dev Biol. 29, 2–16 (2014).

17. A. R. Norman, L. Byrnes, J. F. Reiter, “GC-1 spg cells are most similar to Leydig cells, a testis somatic cell-type, and not germ cells” (preprint, Cell Biology, 2021), doi:10.1101/2021.01.07.425754.

18. M. J. Bitgood, A. P. McMahon, HedgehogandBmpGenes Are Coexpressed at Many Diverse Sites of Cell–Cell Interaction in the Mouse Embryo. Developmental Biology. 172, 126–138 (1995).

19. M. J. Bitgood, L. Shen, A. P. McMahon, Sertoli cell signaling by Desert hedgehog regulates the male germline. Curr Biol. 6, 298–304 (1996).

20. I. Barsoum, H. H. C. Yao, Redundant and differential roles of transcription factors Gli1 and Gli2 in the development of mouse fetal Leydig cells. Biol Reprod. 84, 894–899 (2011).

21. H. H.-C. Yao, W. Whoriskey, B. Capel, Desert Hedgehog/Patched 1 signaling specifies fetal Leydig cell fate in testis organogenesis. Genes Dev. 16, 1433–1440 (2002).

22. A. M. Clark, K. K. Garland, L. D. Russell, Desert hedgehog (Dhh) gene is required in the mouse testis for formation of adult-type Leydig cells and normal development of peritubular cells and seminiferous tubules. Biol. Reprod. 63, 1825–1838 (2000).

23. F. Pierucci-Alves, A. M. Clark, L. D. Russell, A developmental study of the Desert hedgehog-null mouse testis. Biol. Reprod. 65, 1392–1402 (2001).

24. C. B. Bai, W. Auerbach, J. S. Lee, D. Stephen, A. L. Joyner, Gli2, but not Gli1, is required for initial Shh signaling and ectopic activation of the Shh pathway. Development. 129, 4753–4761 (2002).

25. F. He, P. Akbari, R. Mo, J. J. Zhang, C.-C. Hui, P. C. Kim, W. A. Farhat, Adult Gli2+/-;Gli3Δ699/+ Male and Female Mice Display a Spectrum of Genital Malformation. PLoS ONE. 11, e0165958 (2016).

26. K. Lindblad-Toh, M. Garber, O. Zuk, M. F. Lin, B. J. Parker, S. Washietl, P. Kheradpour, J. Ernst, G. Jordan, E. Mauceli, L. D. Ward, C. B. Lowe, A. K. Holloway, M. Clamp, S. Gnerre, J. Alföldi, K. Beal, J. Chang, H. Clawson, J. Cuff, F. Di Palma, S. Fitzgerald, P. Flicek, M. Guttman, M. J. Hubisz, D. B. Jaffe, I. Jungreis, W. J. Kent, D. Kostka, M. Lara, A. L. Martins, T. Massingham, I. Moltke, B. J. Raney, M. D. Rasmussen, J. Robinson, A. Stark, A. J. Vilella, J. Wen, X. Xie, M. C. Zody, Broad Institute Sequencing Platform and Whole Genome Assembly Team, J. Baldwin, T. Bloom, C. W. Chin, D. Heiman, R. Nicol, C. Nusbaum, S. Young, J. Wilkinson, K. C. Worley, C. L. Kovar, D. M. Muzny, R. A. Gibbs, Baylor College of Medicine Human Genome Sequencing Center Sequencing Team, A. Cree, H. H. Dihn, G. Fowler, S. Jhangiani, V. Joshi, S. Lee, L. R. Lewis, L. V. Nazareth, G. Okwuonu, J. Santibanez, W. C. Warren, E. R. Mardis, G. M. Weinstock, R. K. Wilson, Genome Institute at Washington University, K. Delehaunty, D. Dooling, C. Fronik, L. Fulton, B. Fulton, T. Graves, P. Minx, E. Sodergren, E. Birney, E. H. Margulies, J. Herrero, E. D. Green, D. Haussler, A. Siepel, N. Goldman, K. S. Pollard, J. S. Pedersen, E. S. Lander, M. Kellis, A high-resolution map of human evolutionary constraint using 29 mammals. Nature. 478, 476–482 (2011).

27. F. Inoue, N. Ahituv, Decoding enhancers using massively parallel reporter assays. Genomics. 106, 159–164 (2015).

28. M. C. Hofmann, S. Narisawa, R. A. Hess, J. L. Millán, Immortalization of germ cells and somatic testicular cells using the SV40 large T antigen. Exp. Cell Res. 201, 417–435 (1992).

29. F. Inoue, M. Kircher, B. Martin, G. M. Cooper, D. M. Witten, M. T. McManus, N. Ahituv, J. Shendure, A systematic comparison reveals substantial differences in chromosomal versus episomal encoding of enhancer activity. Genome Res. 27, 38–52 (2017).

30. N. Krietenstein, S. Abraham, S. V. Venev, N. Abdennur, J. Gibcus, T.-H. S. Hsieh, K. M. Parsi, L. Yang, R. Maehr, L. A. Mirny, J. Dekker, O. J. Rando, Ultrastructural Details of Mammalian Chromosome Architecture. Mol. Cell (2020), doi:10.1016/j.molcel.2020.03.003.

31. E. K. Unger, K. J. Burke, C. F. Yang, K. J. Bender, P. M. Fuller, N. M. Shah, Medial amygdalar aromatase neurons regulate aggression in both sexes. Cell Rep. 10, 453–462 (2015).

32. K. Hashikawa, Y. Hashikawa, A. Falkner, D. Lin, The neural circuits of mating and fighting in male mice. Curr. Opin. Neurobiol. 38, 27–37 (2016).

33. J. M. Casto, O. B. Ward, A. Bartke, Play, copulation, anatomy, and testosterone in gonadally intact male rats prenatally exposed to flutamide. Physiol. Behav. 79, 633–641 (2003).

34. C. E. Roselli, J. M. Schrunk, H. L. Stadelman, J. A. Resko, F. Stormshak, The effect of aromatase inhibition on the sexual differentiation of the sheep brain. Endocrine. 29, 501–511 (2006).

35. C. H. Phoenix, R. W. Goy, A. A. Gerall, W. C. Young, Organizing action of prenatally administered testosterone propionate on the tissues mediating mating behavior in the female guinea pig. Endocrinology. 65, 369–382 (1959).

36. J. Clarkson, A. E. Herbison, Hypothalamic control of the male neonatal testosterone surge. Phil. Trans. R. Soc. B. 371, 20150115 (2016).

37. K. Raskin, K. de Gendt, A. Duittoz, P. Liere, G. Verhoeven, F. Tronche, S. Mhaouty-Kodja, Conditional inactivation of androgen receptor gene in the nervous system: effects on male behavioral and neuroendocrine responses. J. Neurosci. 29, 4461–4470 (2009).

38. L. A. Lettice, S. J. H. Heaney, L. A. Purdie, L. Li, P. de Beer, B. A. Oostra, D. Goode, G. Elgar, R. E. Hill, E. de Graaff, A long-range Shh enhancer regulates expression in the developing limb and fin and is associated with preaxial polydactyly. Hum. Mol. Genet. 12, 1725–1735 (2003).

39. J. Lopez-Rios, A. Duchesne, D. Speziale, G. Andrey, K. A. Peterson, P. Germann, E. Unal, J. Liu, S. Floriot, S. Barbey, Y. Gallard, M. Müller-Gerbl, A. D. Courtney, C. Klopp, S. Rodriguez, R. Ivanek, C. Beisel, C. Wicking, D. Iber, B. Robert, A. P. McMahon, D. Duboule, R. Zeller, Attenuated sensing of SHH by Ptch1 underlies evolution of bovine limbs. Nature. 511, 46–51 (2014).

40. A. Burga, W. Wang, E. Ben-David, P. C. Wolf, A. M. Ramey, C. Verdugo, K. Lyons, P. G. Parker, L. Kruglyak, A genetic signature of the evolution of loss of flight in the Galapagos cormorant. Science. 356, eaal3345 (2017).

41. M. E. Ritchie, B. Phipson, D. Wu, Y. Hu, C. W. Law, W. Shi, G. K. Smyth, limma powers differential expression analyses for RNA-sequencing and microarray studies. Nucleic Acids Research. 43, e47–e47 (2015).

42. Y. Benjamini, Y. Hochberg, Controlling the False Discovery Rate: A Practical and Powerful Approach to Multiple Testing. Journal of the Royal Statistical Society: Series B (Methodological). 57, 289–300 (1995).

43. K. R. Rosenbloom, C. A. Sloan, V. S. Malladi, T. R. Dreszer, K. Learned, V. M. Kirkup, M. C. Wong, M. Maddren, R. Fang, S. G. Heitner, B. T. Lee, G. P. Barber, R. A. Harte, M. Diekhans, J. C. Long, S. P. Wilder, A. S. Zweig, D. Karolchik, R. M. Kuhn, D. Haussler, W. J. Kent, ENCODE data in the UCSC Genome Browser: year 5 update. Nucleic Acids Res. 41, D56–63 (2013).

44. X. Wang, M. McManus, Lentivirus Production. JoVE, 1499 (2009).

45. F. A. Ran, P. D. Hsu, J. Wright, V. Agarwala, D. A. Scott, F. Zhang, Genome engineering using the CRISPR-Cas9 system. Nat Protoc. 8, 2281–2308 (2013).

46. J. G. Doench, N. Fusi, M. Sullender, M. Hegde, E. W. Vaimberg, K. F. Donovan, I. Smith, Z. Tothova, C. Wilen, R. Orchard, H. W. Virgin, J. Listgarten, D. E. Root, Optimized sgRNA design to maximize activity and minimize off-target effects of CRISPR-Cas9. Nat Biotechnol. 34, 184–191 (2016).

47. E. Dodou, S.-M. Xu, B. L. Black, mef2c is activated directly by myogenic basic helix-loop-helix proteins during skeletal muscle development in vivo. Mechanisms of Development. 120, 1021–1032 (2003).

48. T. D. Schmittgen, K. J. Livak, Analyzing real-time PCR data by the comparative C(T) method. Nat Protoc. 3, 1101–1108 (2008).

49. M. Haeussler, A. S. Zweig, C. Tyner, M. L. Speir, K. R. Rosenbloom, B. J. Raney, C. M. Lee, B. T. Lee, A. S. Hinrichs, J. N. Gonzalez, D. Gibson, M. Diekhans, H. Clawson, J. Casper, G. P. Barber, D. Haussler, R. M. Kuhn, W. J. Kent, The UCSC Genome Browser database: 2019 update. Nucleic Acids Res. 47, D853–D858 (2019).

50. P. H. L. Krijger, G. Geeven, V. Bianchi, C. R. E. Hilvering, W. de Laat, 4C-seq from beginning to end: A detailed protocol for sample preparation and data analysis. Methods. 170, 17–32 (2020).

51. H. Wang, N. Su, L.-C. Wang, X. Wu, S. Bui, K.-J. Chang, A. Nielsen, H.-T. Vo, Y. Luo, X.-J. Ma, in In Situ Hybridization Methods, G. Hauptmann, Ed. (Springer New York, New York, NY, 2015; https://doi.org/10.1007/978-1-4939-2303-8_21), pp. 405–414.

52. T. Ferreira, W. Rasband, ImageJ user guide. ImageJ/Fiji. 1, 155–161 (2012).

53. M. Martinez-Losa, T. E. Tracy, K. Ma, L. Verret, A. Clemente-Perez, A. S. Khan, I. Cobos, K. Ho, L. Gan, L. Mucke, M. Alvarez-Dolado, J. J. Palop, Nav1.1-Overexpressing Interneuron Transplants Restore Brain Rhythms and Cognition in a Mouse Model of Alzheimer’s Disease. Neuron. 98, 75–89.e5 (2018).

54. S. F. Owen, J. D. Berke, A. C. Kreitzer, Fast-Spiking Interneurons Supply Feedforward Control of Bursting, Calcium, and Plasticity for Efficient Learning. Cell. 172, 683–695.e15 (2018).

55. S.-I. Hagiwara, E. Kaushal, S. Paruthiyil, P. J. Pasricha, B. Hasdemir, A. Bhargava, Gastric corticotropin-releasing factor influences mast cell infiltration in a rat model of functional dyspepsia. PLoS ONE. 13, e0203704 (2018).

56. C. Tai, C.-W. Chang, G.-Q. Yu, I. Lopez, X. Yu, X. Wang, W. Guo, L. Mucke, Tau Reduction Prevents Key Features of Autism in Mouse Models. Neuron. 106, 421–437.e11 (2020).

57. K. Scearce-Levie, E. D. Roberson, H. Gerstein, J. A. Cholfin, V. S. Mandiyan, N. M. Shah, J. L. R. Rubenstein, L. Mucke, Abnormal social behaviors in mice lacking Fgf17. Genes Brain Behav. 7, 344–354 (2008).

58. G. Krabbe, S. S. Minami, J. I. Etchegaray, P. Taneja, B. Djukic, D. Davalos, D. Le, I. Lo, L. Zhan, M. C. Reichert, F. Sayed, M. Merlini, M. E. Ward, D. C. Perry, S. E. Lee, A. Sias, C. N. Parkhurst, W.-B. Gan, K. Akassoglou, B. L. Miller, R. V. Farese, L. Gan, Microglial NFκB-TNFα hyperactivation induces obsessive-compulsive behavior in mouse models of progranulin-deficient frontotemporal dementia. Proc. Natl. Acad. Sci. U.S.A. 114, 5029–5034 (2017).

59. H. Belinson, J. Nakatani, B. A. Babineau, R. Y. Birnbaum, J. Ellegood, M. Bershteyn, R. J. McEvilly, J. M. Long, K. Willert, O. D. Klein, N. Ahituv, J. P. Lerch, M. G. Rosenfeld, A. Wynshaw-Boris, Prenatal β-catenin/Brn2/Tbr2 transcriptional cascade regulates adult social and stereotypic behaviors. Mol. Psychiatry. 21, 1417–1433 (2016).

60. C. C. Cheung, W. C. Krause, R. H. Edwards, C. F. Yang, N. M. Shah, T. S. Hnasko, H. A. Ingraham, Sex-dependent changes in metabolism and behavior, as well as reduced anxiety after eliminating ventromedial hypothalamus excitatory output. Mol Metab. 4, 857–866 (2015).

61. L. Wang, J. Simms, C. J. Peters, M. Tynan-La Fontaine, K. Li, T. M. Gill, Y. N. Jan, L. Y. Jan, TMEM16B Calcium-Activated Chloride Channels Regulate Action Potential Firing in Lateral Septum and Aggression in Male Mice. J. Neurosci. 39, 7102–7117 (2019).

62. Z. Sahin, A. Szczepny, E. A. McLaughlin, M. L. Meistrich, W. Zhou, I. Ustunel, K. L. Loveland, Dynamic Hedgehog signalling pathway activity in germline stem cells. Andrology. 2, 267–274 (2014).

63. G. La Sala, D. Marazziti, C. Di Pietro, E. Golini, R. Matteoni, G. P. Tocchini-Valentini, Modulation of Dhh signaling and altered Sertoli cell function in mice lacking the GPR37-prosaposin receptor. FASEB J. 29, 2059–2069 (2015).

64. H. Kim, H. Xu, Q. Yao, W. Li, Q. Huang, P. Outeda, V. Cebotaru, M. Chiaravalli, A. Boletta, K. Piontek, G. G. Germino, E. J. Weinman, T. Watnick, F. Qian, Ciliary membrane proteins traffic through the Golgi via a Rabep1/GGA1/Arl3-dependent mechanism. Nat Commun. 5, 5482 (2014).

65. R. V. Mettus, S. G. Rane, Characterization of the abnormal pancreatic development, reduced growth and infertility in Cdk4 mutant mice. Oncogene. 22, 8413–8421 (2003).

66. J. A. Costoya, R. M. Hobbs, M. Barna, G. Cattoretti, K. Manova, M. Sukhwani, K. E. Orwig, D. J. Wolgemuth, P. P. Pandolfi, Essential role of Plzf in maintenance of spermatogonial stem cells. Nat. Genet. 36, 653–659 (2004).

67. J. Qin, M.-J. Tsai, S. Y. Tsai, Essential roles of COUP-TFII in Leydig cell differentiation and male fertility. PLoS ONE. 3, e3285 (2008).

68. R. Kimura, K. Yoshizaki, N. Osumi, Dynamic expression patterns of Pax6 during spermatogenesis in the mouse. J. Anat. 227, 1–9 (2015).

